# Designing monomeric IFNγ: the significance of domain-swapped dimer structure in IFNγ immune responses

**DOI:** 10.1101/2024.02.18.580912

**Authors:** Yota Goto, Takamitsu Miyafusa, Shinya Honda

## Abstract

IFNγ can initiate immune responses by inducing the expression of major histocompatibility complex molecules, suggesting its potential for cancer immunotherapy. However, it also has an immunosuppressive function that limits its application as a therapeutic agent. IFNγ has a characteristic domain-swapped dimer structure with two of the six α-helices exchanged with each other. As we hypothesized that the contrasting functions of IFNγ could be attributed to its unique domain-swapped structure, we designed monomeric IFNγ by transforming the domain-swapped dimer structure of wild-type IFNγ. We conjectured the evolution of this domain-swapped dimer and hypothesized that the current IFNγ structure emerged through shortening of the loop structure at the base of the swapped domain and the accumulation of hydrophobic amino acids at the newly generated interface during domain-swapping. We then designed and generated a stable monomeric IFNγ by retracing this evolutionary process, complementing the lost loop structure with a linker and replacing the accumulated hydrophobic amino acids with hydrophilic ones. We determined that the designed variant was a monomer based on molecular size and number of epitopes and exhibited activity in cell-based assays. Notably, the monomeric IFNγ showed a qualitatively similar balance between immunostimulatory and immunosuppressive gene expression as wild-type IFNγ. This study demonstrates that the structural format of IFNγ affects the strength of its activity rather than regulating the fate of downstream gene expression.

## INTRODUCTION

Interferons constitute a family of proteins that are produced by leukocytes and possess antiviral activity.^1^ Among these, IFNγ is the only type II interferon^2^. It is distinct from type I interferons, such as IFNα and IFNβ, or the more recently discovered type III interferons, which include IFNλ, in terms of sequence homology and target receptors on the cell surface.^3,4^ While all types of interferons regulate diverse immunoregulatory functions, IFNγ is particularly noted for its ability to upregulate the expression of both major histocompatibility complex (MHC) class I and II molecules as well as for its potent antiproliferative effects.^5–7^ One of its major anti-tumor effects is the promotion of antigen presentation via MHC upregulation.^8^

Having been the subject of multiple clinical trials, IFNγ is not yet approved as an immuno-oncology drug.^9^ One aspect limiting its clinical application is the difficulty in controlling its multiple functions. For instance, IFNγ can activate immunity by upregulating the expression of MHC molecules, yet it can also suppress immunity by upregulating the expression of programmed death-ligand 1 (PD-L1). These contrasting functions complicate its use for cancer immunotherapy, with some clinical research showing that IFNγ enhances tumor growth instead of inhibiting it.^10,11^

IFNγ has a domain-swapped dimer structure with two of the six α-helices exchanged with each other.^12,13^ This homodimer forms a 2:2:2 complex with two receptors, IFNγR1 and IFNγR2, on the cell surface, thus initiating signal transduction.^14–16^ Both receptors belong to the class 2 cytokine receptor superfamily, which includes receptors to IFNα and interleukin (IL)-10.^17^ The following pathway is considered one of the primary mechanisms through which IFNγ stimulation drives gene expression. First, the IFNγ/IFNγR1/IFNγR2 ternary complex undergoes endocytosis into the cytoplasm. Thereafter, JAK1 and JAK2, which associate with the intracellular domains of IFNγR1 and IFNγR2, are activated and subsequently phosphorylate Y701 in STAT1. The phosphorylated STAT1 (pSTAT1) subsequently dimerizes, forms a complex with IFNγ and IFNγR1, and then translocates into the nucleus to upregulate target gene expression.^18–22^ While unphosphorylated STAT1 migrates into the nucleus via a carrier-free mechanism,^23–25^ nuclear translation of the pSTAT1 complex occurs via the Ran/importin pathway, which requires a nuclear localization sequence (NLS). According to studies by Subramaniam et al., the NLS at the C-terminus of IFNγ (^126^KTGKRKR^132^) is required for the nuclear localization of the pSTAT1 complex. In addition, this sequence binds to the intracellular region of IFNγR1 and triggers phosphorylation of STAT1, an event implicated in the antiviral activity of IFNγ.^26–30^

Despite extensive research into IFNγ signal transduction, the mechanisms underlying its diverse functions are not fully understood, and the associated regulatory mechanisms have not been established. Regulating IFNγ function could prove valuable for cancer immunotherapy and the treatment of autoimmune diseases. One strategy to achieve this is through the design of IFNγ variants with controllable activity.

In this study, we aimed to design monomeric IFNγ by transforming its domain-swapped dimeric structure into a monomer. One of the IFNγ variants reported to date is GIFN2, in which only one subunit of dimeric IFNγ is altered and its receptor-binding activity is eliminated.^16^ This variant was found to retain the induction of MHC gene expression while decreasing the induction of PD-L1 gene expression. Therefore, this variant has a reduced immunosuppressive function while maintaining its immunostimulatory function. Structural alterations of IFNγ therefore enable us to create variants that retain only some IFNγ functions. GIFN2 has the same domain-swapped dimer structure as wild-type IFNγ (WT), although some amino acids in the receptor-binding regions are altered. In contrast, monomeric IFNγ, which is the subject of this study, structurally differs from GIFN2, as it lacks one of the two subunits, which potentially gives rise to functional alterations. The structure of the current domain-swapped IFNγ is believed to have evolved from a monomeric form to acquire its current functions.^31^ Therefore, in designing the monomeric IFNγ, we aimed to trace back a hypothetical scenario of IFNγ evolution. The engineered variant proved to be a monomer and displayed an activity comparable to that of WT. However, it did not result in the anticipated simplification of functionality. This discovery unveiled the intricate nature of the structure-function relationship of IFNγ, highlighting that the intracellular signal intensity of the ligand, rather than its structure, is responsible in IFNγ immune responses. These findings will provide new guidelines for regulating the diverse functions of IFNγ.

## RESULTS

As an antecedent to designing a monomeric form of IFNγ, we deduced the origin of its unique structure, a domain-swapped dimer, as illustrated in Figure 1. Domain-swapped dimers are characterized by the exchange of certain domains (swapped domains). This homodimer consisted of two open monomers. It is highly unlikely that such a unique structure would have arisen abruptly during molecular evolution. Instead, a simpler structure plausibly served as the starting point for a gradual increase in complexity. Specifically, the formation of domain-swapped dimers involves the initial appearance of a closed monomer, in which the swapped domain is arranged within the same molecule. Subsequently, shortening of the loop at the base of the swapped domain (hinge loop) occurs, resulting in the formation of an open monomer and, ultimately, a domain-swapped dimer. Although it may be difficult to validate this hypothetical scenario of molecular evolution, it has been put forth in several studies.^31,32^

**Figure 1.**
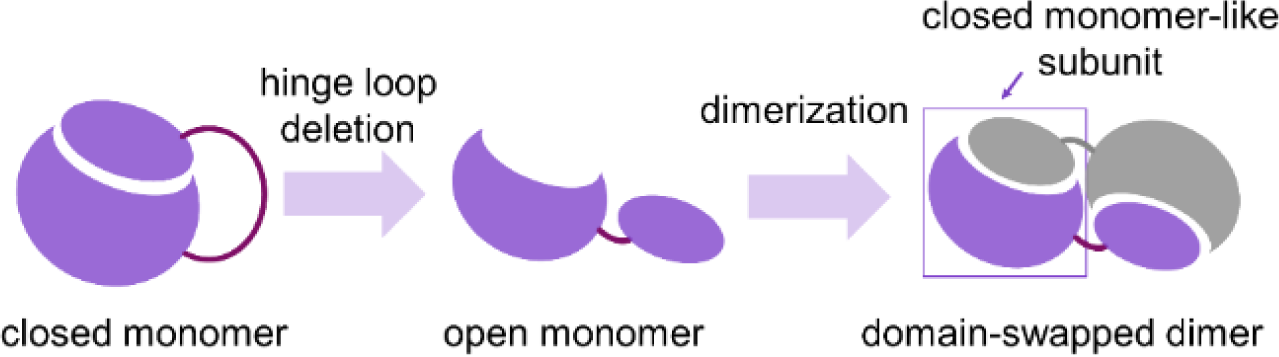
Hypothetical molecular evolution of domain-swapped dimers. It is believed that the open monomer evolved from the closed monomer through shortening of the hinge loop and that the formation of a domain-swapped dimer is completed by the dimerization of the open monomers. Although the domain-swapped dimer is composed of two open monomers, it can also be described as comprising two closed monomer-like subunits.

IFNγ is a homodimer composed of two open monomers. However, it can also be viewed as a pseudo-dimer structure with two closed monomer-like subunits, each consisting of two polypeptides. Thus, the domain-swapped dimer has two interfaces. One is between the open monomers (domain interface), and the other is between two closed monomer-like subunits (subunit interface). In addition to the domain interface, interactions at the subunit interface are also considered as factors that stabilize the domain-swapped dimer.

IFNγ has a relatively large subunit interface in comparison to domain-swapped dimers with small subunit interfaces, such as IL5 and IL10 (Figure 2A).^33,34^ In immature domain-swapped dimers formed via the association of open monomers, the corresponding surface of the subunit interface is hydrophilic, and the subunit interface is expected to be small. Based on these observations, it can be inferred that, during evolution, IFNγ acquired a large subunit interface through the substitution and accumulation of hydrophobic amino acids, starting from structures with small subunit interfaces, similar to that of IL5 and IL10. The hypothesized molecular evolution scenario rationalized the design of monomeric IFNγ.

**Figure 2.**
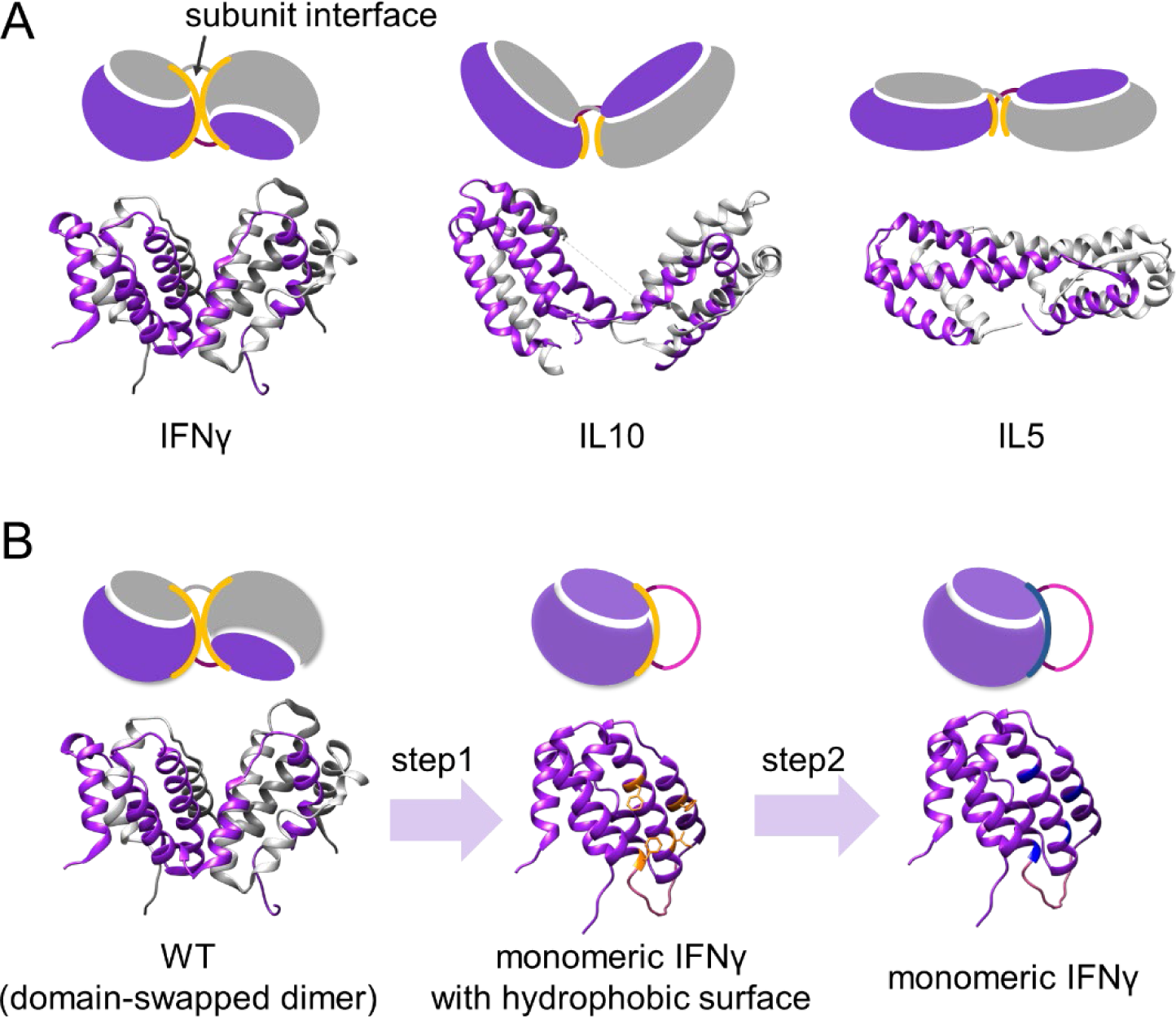
Proteins with domain-swapped dimers and the approach to monomerization. (A) Structures of IFNγ (PDB: 1FG9), IL10 (PDB: 6X93), and IL5 (PDB: 1HUL). Monomerized variants of IL10 and IL5 engineered via linker insertion have been previously reported. ^35–37^ (B) Two-step approach for designing monomeric IFNγ. Step 1: A swapped domain can be structurally rearranged from an open monomer form to a closed monomer structure via linker insertion. Hydrophobic amino acids in the subunit interface are depicted in yellow. Step 2: By replacing the hydrophobic amino acids (yellow) with hydrophilic amino acids (blue), the solubility of the monomer will be increased.

Monomeric IFNγ was generated in the following two-step approach (Figure 2B). First, a linker was inserted into the hinge loop (step 1). This allowed the adoption of a closed monomer corresponding to the precursor before shortening of the hinge loop. Monomerized variants of IL5 and IL10 have been obtained by inserting a linker into the hinge loop, indicating that the linker insertion method is generally applicable.^35–37^ However, in contrast to IL5 and IL10, IFNγ has a large subunit interface, and it is expected that hydrophobic amino acids in the subunit interface may be exposed to the solvent and destabilize the structure when adopting a closed monomer structure. Hence, these amino acids were replaced to render the surface hydrophilic (step 2). We presume that this two-step approach is necessary for obtaining a stable monomer.

### Linker insertion (Step 1)

For linker insertion, the GS linker was chosen as it is a flexible linker. Structural prediction was performed using Alphafold2^38^ to calculate the appropriate linker length that would allow the swapped domain to be arranged within the same molecule. The results suggested that a linker of at least six residues was required for closed monomer formation. Therefore, we decided to use a nine-residue linker, GGGGSGGGG, which was slightly longer than the predicted minimum linker length, and inserted it after residue 86N of the hinge loop to generate a linker-inserted variant (mIFNG1).

Both WT and mIFNG1 proteins were expressed in *Escherichia coli* and obtained as inclusion bodies. However, aggregation occurred during refolding via dialysis under neutral conditions in Tris-HCl buffer (pH 8.0). mIFNG1 showed particularly strong aggregation and could not be prepared in its soluble form. Considering the high pI value of IFNγ (calc. pI: 9.4), we attempted refolding via tangential flow filtration (TFF) under acidic conditions in sodium acetate buffer (pH 5.0) to minimize aggregation. Consequently, WT and its variants were obtained without aggregation. Thus, the refolded WT and its variants were subjected to subsequent analyses under acidic conditions.

The molecular size (molecular weight [MW] and sedimentation coefficient [*s20,w*]) of WT and mIFNG1 was verified using analytical ultracentrifugation (AUC). There was good agreement between the theoretical and observed molecular sizes for the WT, whereas mIFNG1 was found to be in soluble aggregates 10–30 times larger than the theoretical molecular size of a monomer (Figure 3A, B, Table S1).

**Figure 3.**
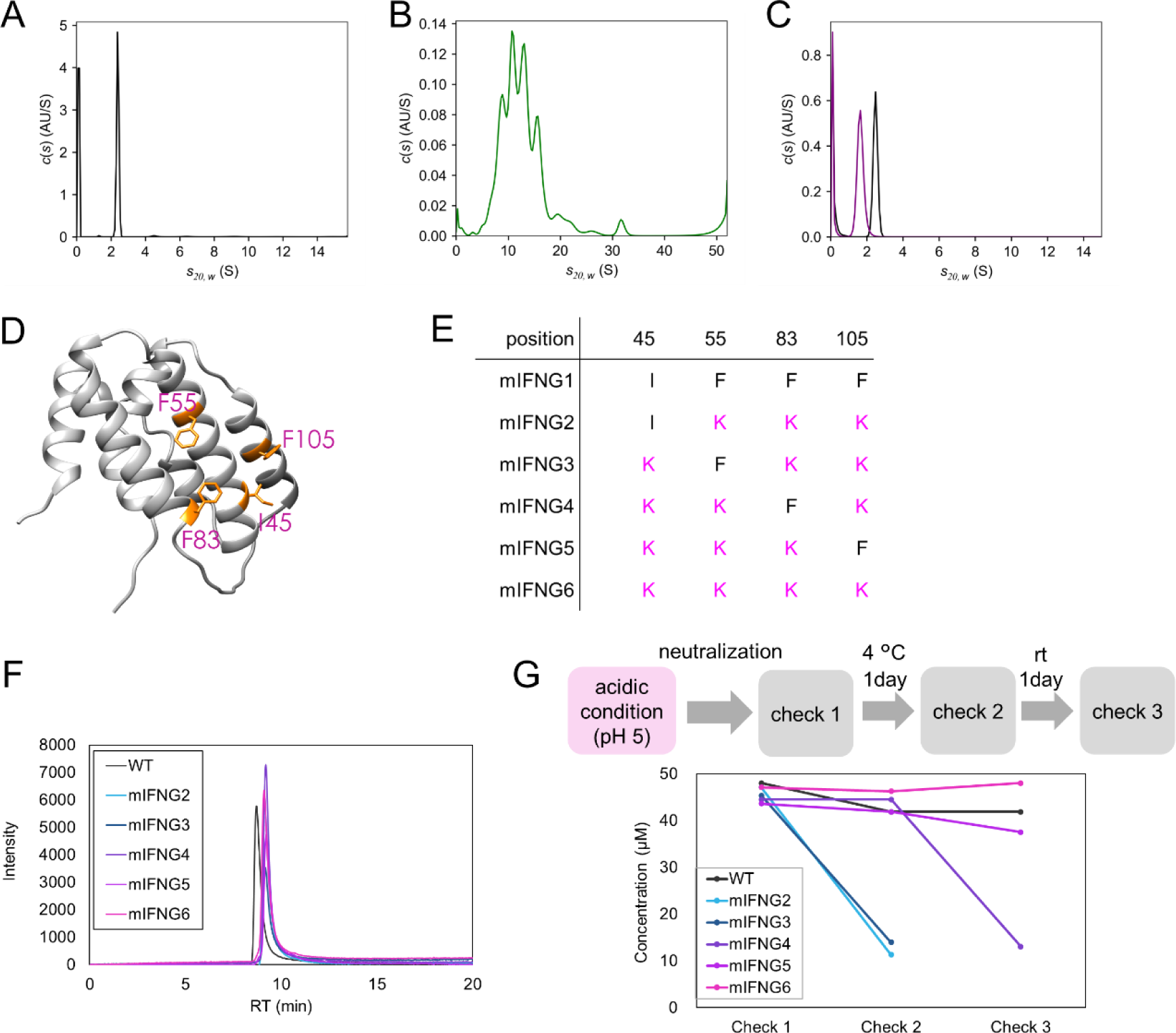
Characterization of IFNγ variants. (A) Sedimentation coefficient distribution of WT measured at 20 °C and pH 5.0. (B) Sedimentation coefficient distribution of mIFNG1 measured at 20 °C and pH 5.0. (C) Sedimentation coefficient distributions of WT (black) and mIFNG5 (purple) measured at 10 °C and pH 5.0. (D) Four hydrophobic amino acids (yellow) examined in the subunit interface. (E) Names of the variants in which the hydrophobic amino acid was replaced with lysine. (F) Analytical SEC results of variants using HPLC measured at pH 5.0. (G) Evaluation of the aggregation propensity of variants under neutral conditions. The concentration of the sample in supernatant was calculated from the absorbance (A280) at every checkpoint.

### Hydrophobic amino acid substitution at the subunit interface (Step 2)

To replace the hydrophobic amino acids at the subunit interface, we focused on four amino acids of mIFNG1, namely, I45, F55, F83, and F105 (Figure 3D). Using the same expression and purification method as for mIFNG1, several variants were generated in which these amino acids were replaced with the hydrophilic amino acid lysine (K) (Figure 3E). The approximate molecular sizes of these variants were estimated from their retention times using size exclusion chromatography (SEC) during the final purification step. Compared to the aggregated variant of mIFNG1, variants with three or four hydrophobic amino acids replaced by lysine were eluted at later retention times, suggesting that these existed as monomers rather than aggregates. To analyze the molecular size of these variants in detail, analytical SEC was performed using HPLC, which has a higher resolution. All variants with three or more substitutions were observed as a single peak with a molecular weight smaller than that of WT (Figure 3F). Therefore, these variants were expected to be monomers.

The variants dissolved well in acidic conditions. Therefore, for further functional evaluation, the aggregation propensity of these variants was investigated under neutral conditions, according to the scheme shown in Figure 3G. mIFNG2 and mIFNG3 showed considerable aggregation at Check 2. mIFNG4 showed no aggregation at Check 2 but aggregated at Check 3. In contrast, WT, mIFNG5, and mIFNG6 did not show apparent aggregates even at Check3. From this assay, we estimated the contribution of the four hydrophobic amino acids of interest to the degree of aggregation, assuming that the hydrophobic amino acids in the variants with a later onset of aggregation contributed less. The order of the aggregation contribution was as follows: F55 = I45 > F83 > F105.

Based on these results, mIFNG5, a triple mutant (I45K/F55K/F83K), was determined to be the least modified variant with the lowest aggregation propensity. Therefore, mIFNG5 was used for further evaluation to determine whether this variant was monomeric and its biological properties.

### Verification of a monomeric IFNγ variant

The molecular sizes (MW and *s20,w*) of WT and mIFNG5 were compared using AUC, as mentioned above. There was good agreement between the theoretical molecular size of the monomer and the observed size for mIFNG5 (Figure 3C, Table S1).

Next, the number of epitopes within the molecule was measured by performing an epitope binning assay under neutral conditions using bio-layer interferometry (BLI).^39^ In this assay, anti-IFNγ antibody was immobilized on a sensor, and measurements were performed by binding WT or mIFNG5, followed by addition of the same antibody to determine the number of epitopes. Both WT and mIFNG5 bound to the antibody-immobilized sensor, as shown in Figure 4A. Furthermore, WT could bind to the added antibody while bound to the antibody on the sensor. In contrast, mIFNG5 did not bind to the added antibody in its bound state. These results indicate that WT possesses two identical epitopes owing to its domain-swapped dimeric structure, whereas mIFNG5 has one epitope owing to its monomeric structure.

**Figure 4.**
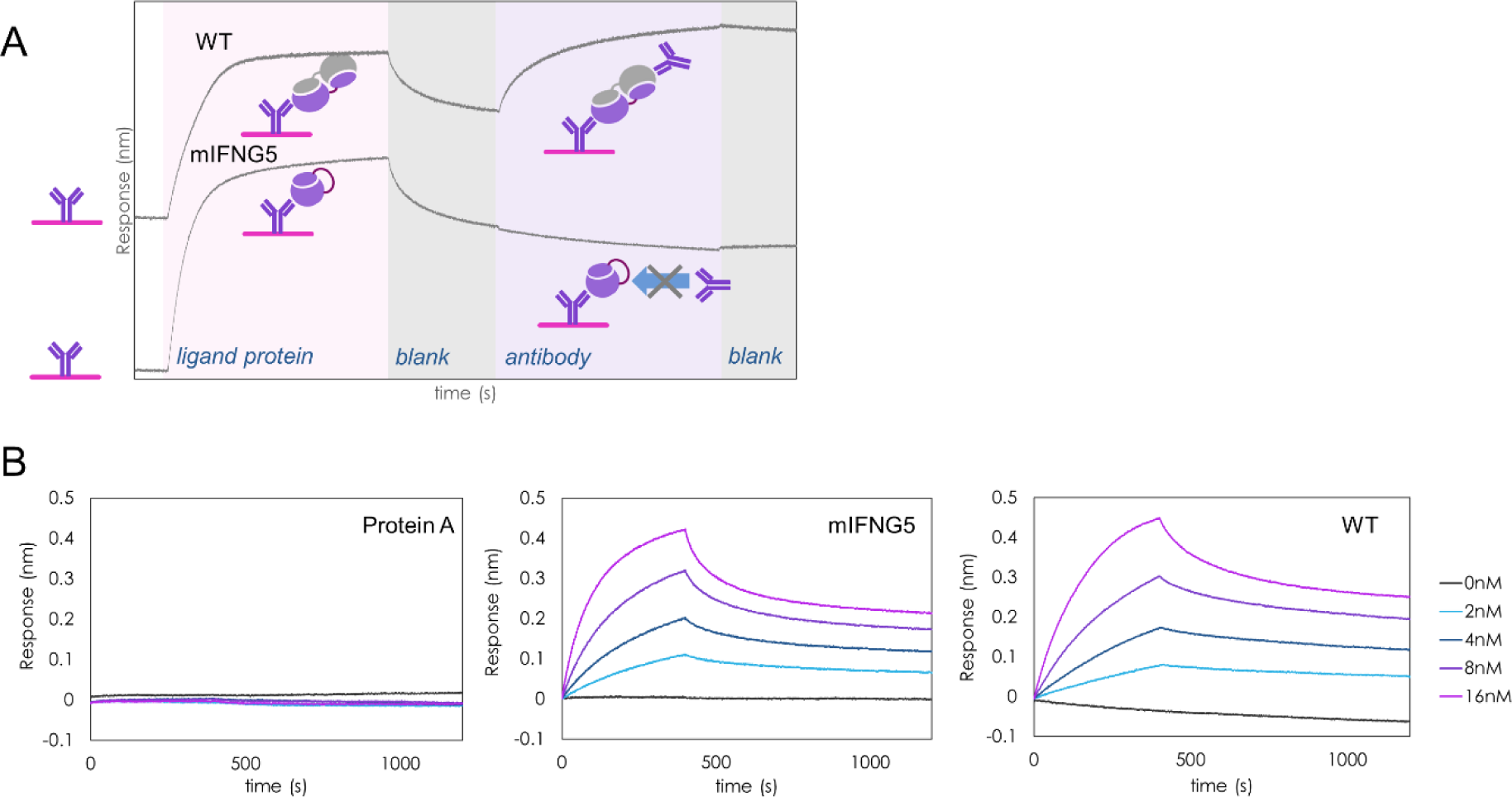
BLI binding assays of monomeric IFNγ. (A) Epitope binning assay of WT and mIFNG5. Biosensors immobilized with anti-IFNγ antibody were immersed into WT- or mIFNG5-containing solutions. Subsequently, the sensors were immersed in the same anti-IFNγ antibody solution. Diagrams illustrate the possible binding of analytes to anti-IFNγ antibody. (B) Representative sensorgrams of BLI measurements examining the interactions of IFNγR1 with protein A (left; control), mIFNG5 (center), and WT (right). Biosensors immobilized with IFNγR1 were immersed into solutions containing a series of analyte concentrations.

### Receptor binding evaluation

To evaluate receptor binding, a kinetic assay of IFNγR1 with WT and mIFNG5 was performed using BLI, and the result was compared to the reported values of a kinetic assay using surface plasmon resonance^40^ (Table S2). mIFNG5 showed typical sensorgrams, as did the WT, whereas the negative control protein A showed no signal, indicating that both proteins specifically bind to IFNγR1 immobilized on the sensor (Figure 4B). The equilibrium dissociation constant (*K*_d_) of mIFNG5 was comparable to that of WT (1/3 of WT), confirming that this variant interacts with IFNγR1.

### Analysis for upstream signaling

IFNγ is internalized via endocytosis after binding to IFNγR1 and IFNγR2. Subsequently, the intracellular domain of IFNγR1 binds to IFNγ in the cytoplasm. This leads to the phosphorylation of IFNγR1 and the subsequent phosphorylation of STAT1. Therefore, we measured the phosphorylation of STAT1, which is an upstream indicator of signal transduction, to assess signal transduction. We thought it would be useful to examine not only the comparison between the wild-type and monomeric form but also the presence or absence of an NLS, which is known to play an influential role in the activity of IFNγ.^27^ Therefore, in addition to WT and mIFNG5, NLS (^126^KTGKRKR^132^)-deleted variants of WT (IFNG(NLS^-^)) and mIFNG5 (mIFNG5(NLS^-^)) were generated and analyzed for their activities in cells. Detection of pSTAT1 via AlphaLISA using HeLa cells showed that the EC_50_ and *E*_max_ values of mIFNG5 were approximately 1/500 and 60%, respectively, relative to those of WT (Figure 5A). This indicates that mIFNG5 maintains STAT1 phosphorylation activity, although it is weaker than that of the WT. The EC_50_ and *E*_max_ values of IFNG(NLS^-^) were 1/100 and 60%, respectively, compared with those of the WT. Little signal was observed for mIFNG1 and mIFNG5(NLS^-^) (Figure 5B, C).

**Figure 5.**
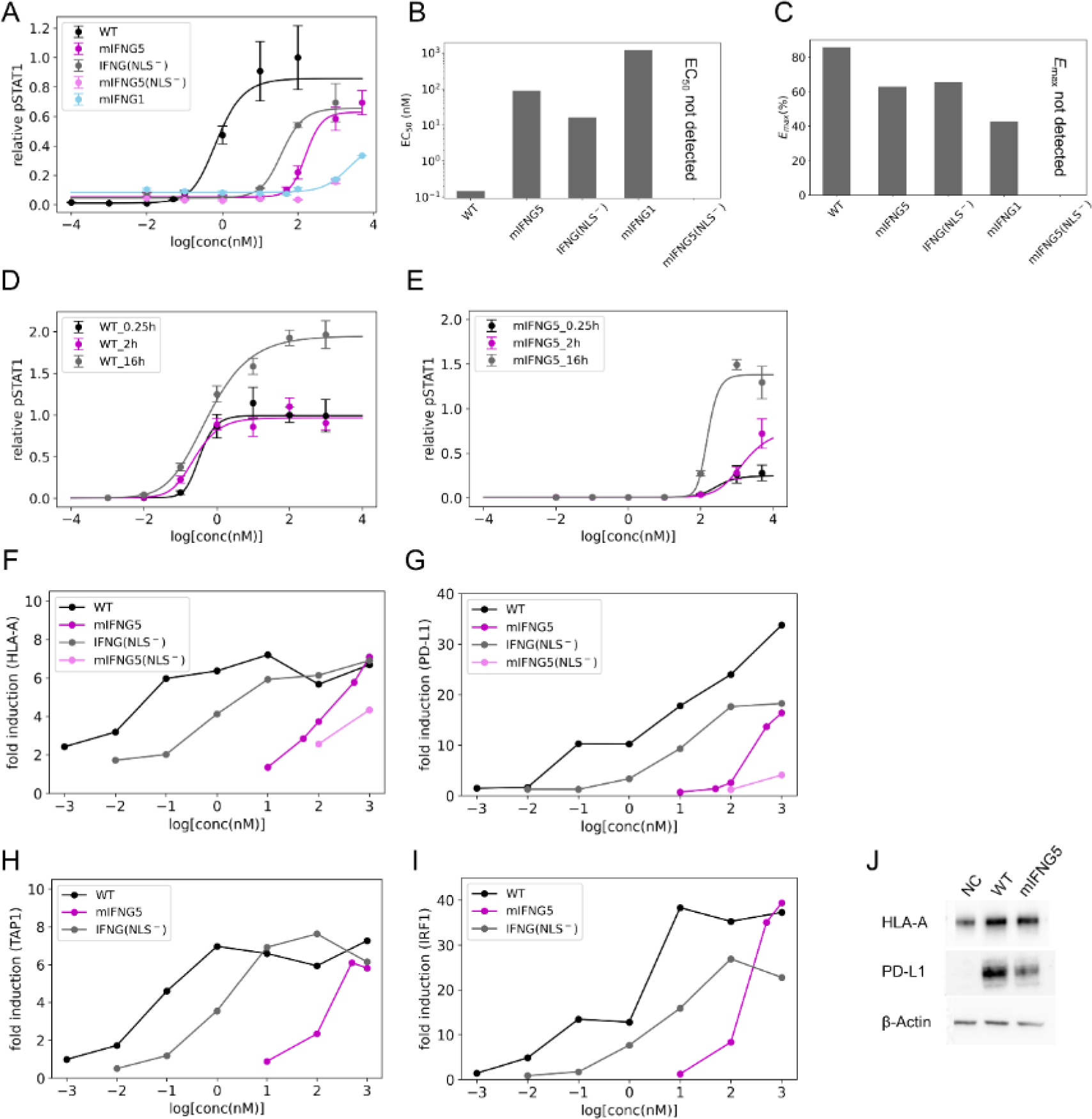
Upstream and downstream signaling pathway analyses upon the stimulation of IFNγ variants. (A) STAT1 phosphorylation activity assay. HeLa cells were treated with WT or variants for 0.25 h. Phosphorylated STAT1 (pSTAT1) was measured using AlphaLISA. Data represent the mean ± standard error of the mean (SEM) of n=3 biological replicates, except for mIFNG1. (Data for mIFNG1 represent the mean ± SEM of n=2 biological replicates.) Solid lines were calculated via 4-parameter logistic curve fitting. (B) EC_50_ (nM) of STAT1 phosphorylation by the variants obtained from the fitting calculation. (C) Relative *E*_max_ (%) of STAT1 phosphorylation by the variants. The values were normalized to measurements at 100 nM WT stimulation. (D, E) Time dependence of STAT1 phosphorylation by WT (D) and mIFNG5 (E). HeLa cells were treated with WT or mIFNG5 for 0.25, 2, or 16 h. Phosphorylated STAT1 (pSTAT1) was measured using AlphaLISA. Data represent the mean ± standard error of the mean (SEM) of n=3 biological replicates. (F, G, H, I) Fold induction of target genes HLA-A (F), PD-L1 (G), TAP1 (H), and IRF1 (I) compared to the unstimulated state (NC). HeLa cells were treated with WT and variants for 48 h, whereafter mRNA was extracted from each sample. Gene expression levels of the target genes and the reference 18S rRNA were quantified based on the amount of corresponding mRNA determined via RT-qPCR. Data represent values obtained from a single biological replicate (n=1). (J) Western blotting analysis of HLA-A, PD-L1, and β-actin in cell lysates. HeLa cells were treated with 1000 nM of WT or 1000 nM of mIFNG5 for 48 h, and the cell lysates were analyzed.

Next, the kinetics of STAT1 phosphorylation were further investigated. STAT1 phosphorylation is an essential step in IFNγ signaling, as pSTAT1 serves as a transcription factor, inducing the expression of various genes, including itself.^41^ If IFNγ is still available and active at the time of the induction of new STAT1 synthesis, further phosphorylation occurs, resulting in signal enhancement. Therefore, pSTAT1 was measured at 0.25, 2, and 16 h after stimulation with WT and mIFNG5 (Figure 5D, E). pSTAT1 levels were similar at 0.25 and 2 h after stimulation with the WT, but a significant increase was observed at 16 h. mIFNG5 induced a slight increase in pSTAT1 at 2 h compared to 0.25 h and a significant increase at 16 h, as in the case of WT. This difference may indicate that STAT1 phosphorylation induced by mIFNG5 is slower than that triggered by WT owing to its monomeric nature. However, even in the case of mIFNG5, pSTAT1 levels were further enhanced at 16 h, as with the WT, indicating that STAT1 expression is also induced by mIFNG5 and that the induced STAT1 is further phosphorylated.

### Analysis for downstream signaling

Gene expressions induced by IFNγ signaling were analyzed to evaluate downstream signaling. HLA-A, an MHC class I molecule associated with immune activation, and PD-L1, a gene associated with immunosuppression, were selected as the target genes for analysis. In addition, TAP1, which is responsible for transporting antigens to MHC class I molecules, and IRF1, a gene implicated in PD-L1 expression, were analyzed as immune activation- and immunosuppression-associated genes, respectively. To quantify gene expression, relative quantification of transcript levels was performed using quantitative reverse transcription PCR (RT-qPCR). By treatment with different variants and their concentrations, we compared the immune responses via IFNγ signaling (Figure 5F–I, S1). In addition to RT-qPCR, protein expression of the target genes was assessed via Western blotting (Figure 5J).

In WT, HLA-A expression reached a plateau level that was approximately 6–7 times higher than that in the unstimulated state, even at a low concentration (0.1 nM), whereas PD-L1 expression increased proportionally up to 1000 nM, indicating a different concentration-dependent response to IFNγ stimulation. For mIFNG5, stimulation at 500 nM resulted in gene expression equivalent to that of WT stimulated at 0.1 nM. At low concentrations, only HLA-A expression was induced, whereas PD-L1 expression remained low. In contrast, at high concentrations, the induction of both HLA-A and PD-L1 expression was observed, as in the WT. IFNG(NLS^-^) exhibited a mid-level gene expression-inducing activity, with 10 nM-induced stimulation corresponding to that induced by 0.1 nM of WT (Figure 5F, G). Therefore, mIFNG5 and IFNG(NLS^-^) exhibited 1/5000 and 1/100 of the expression-inducing activity, respectively, when compared to the WT. TAP1 and IRF1 showed a similar dependence on ligand concentration as HLA-A and PD-L1, respectively. Changes in gene expression in response to the IFNγ variants were also comparable between HLA-A and TAP1, as well as between PD-L1 and IRF1 (Figure 5H, I).

## DISCUSSION

### Design of monomeric IFNγ through a two-step approach

In this study, we sought to obtain monomeric IFNγ through the transformation of a domain-swapped dimer into a monomer. First, we found that the linker-inserted variant mIFNG1, obtained as a soluble protein, was present in the form of aggregates of 10–30 molecules. This observation was expected in the light of the highly hydrophobic subunit interface of IFNγ being exposed to the solvent by the linker insertion. Subsequently, we found that replacing three or four hydrophobic amino acids at the subunit interface with lysine resulted in monomer formation. mIFNG5 was confirmed as monomeric in terms of both molecular size and number of epitopes. The contribution of hydrophobic amino acids was evaluated by examining the aggregation properties of variants, which showed that I45 and F55 had a greater impact on aggregation. Notably, both I45 and F55 are in the third α-helix from the N-terminus, which is located at the center of the molecule in the domain-swapped dimer. Interestingly, the amino acids in the most hydrophobic environment had a more pronounced effect on aggregation after monomerization.

Based on the observation that mIFNG1 exists as an aggregate, the evolutionary origins of IFNγ can be speculated. Linker insertion into the hinge loop allowed the formation of a closed monomer, although the linker-inserted variant also had the potential to form an open monomer. Essentially, mIFNG1 would exist as a domain-swapped dimer if the hydrophobic subunit interface was positioned within the molecule, similar to that in WT. However, no domain-swapped dimers were observed. This suggests that intramolecular arrangement into a closed monomer occurs at a faster rate than intermolecular domain swapping. Consequently, mIFNG1 adopts a monomeric structure, in which the hydrophobic subunit interface is exposed to the solvent, leading to its aggregation. This implied that the hinge loop must initially be shortened to form a domain-swapped dimer from the monomer, thereby preventing it from becoming a closed monomer. Hydrophobic amino acids then accumulate at the subunit interface, enlarging the interface. This finding supports our hypothesis that IFNγ evolved from a closed monomer through a domain-swapped dimer with a small subunit interface, like IL5 or IL10, into its current state as a domain-swapped dimer with a large subunit interface.

To further discuss evolutionary origins, a phylogenetic tree was constructed for IFNγ in various species based on sequence homology. The crystal structures of each IFNγ and the predicted structure generated by Alphafold2 are also shown in the tree (Figure S2). All IFNγs analyzed here represented domain-swapped dimers. Among them, three mammalian IFNγs (human, rabbit, and bovine) were structurally similar and had a large subunit interface. In contrast, the other IFNγs had a small subunit interface. The result showing the domain-swapped dimer structure with a small subunit interface in distant IFNγ species also supports the above-described hypothesis.

In this study, we focused on the four hydrophobic amino acids I45, F55, F83, and F93 (F105 in mIFNG1) present in the subunit interface of human IFNγ. Therefore, the amino acid sequences of several IFNγs with a large subunit interface (i.e., mammalian IFNγ) were compared (Figure S3). The amino acid residues corresponding to positions I45, F55, and F93 were conserved among the three species. The residues corresponding to the F83 position are phenylalanine (F) and leucine (L) in rabbit and bovine IFNγ, respectively. All these residues are hydrophobic. Therefore, the hydrophobic nature of these residues is crucial for the formation of a domain-swapped dimer with a large subunit interface.

### Activity of monomeric IFNγ and the significance of a domain-swapped dimer

A kinetic assay using BLI revealed that mIFNG5, a monomeric IFNγ, maintains its affinity for IFNγR1. Additionally, pSTAT1 assays demonstrated that both mIFNG5 and IFNG(NLS^-^) retained STAT1 phosphorylation activity, similar to WT. These findings unequivocally establish that neither the structural format, i.e., domain-swapped dimer or monomer, nor the presence or absence of an NLS are essential for IFNγ activity.

The structural format of the ligands did, however, influence the magnitude of activity. First, IFNG(NLS^-^) exhibited activity (EC_50_) that was two orders of magnitude weaker compared to that of WT in pSTAT1 assays, which is consistent with a previous study^26^ that reported the reduced activity upon NLS deletion, highlighting the importance of NLS not only in nuclear localization but also in STAT1 phosphorylation.^23^ Second, mIFNG5 displayed an EC_50_ two orders of magnitude lower than that of WT, suggesting that the difference in activity can be attributed to the structural format of the domain-swapped dimer and monomer. The underlying reasons for this distinction remain unclear. However, potential factors contributing to this distinction may include improved recruitment efficiency of the receptor complex, increased efficiency of endocytosis after receptor binding, enhanced dimerization efficiency of pSTAT1, and increased molecular stability. The kinetics of STAT1 phosphorylation indicate that mIFNG5, like the WT, remains active for at least 16 h in the culture medium. Therefore, the difference in stability between the dimeric and monomeric structures would not be the only reason for the low activity of mIFNG5. In any case, WT maintained its high activity by presenting a unique structure. The domain-swapped dimer is important for the completion of this structural format.

### Signal intensity-dependent gene expression

The expression balance between HLA-A and PD-L1 varied depending on ligand concentration. IFNγ variants act as immunostimulatory factors in a certain concentration range, while exhibiting immunosuppressive effects in another. For example, at concentrations up to 100 nM, mIFNG5 upregulated HLA-A expression and minimally induced PD-L1 expression, which would result in enhanced immune activation. In contrast, at other concentrations (100 nM or higher), PD-L1 was induced to a comparable extent relative to that induced by WT, which would result in immune suppression. A similar gene expression balance was observed for IFNG(NLS^-^).

The correlation between HLA-A and PD-L1 expression levels is shown in Figure 6. Notably, the ratio of HLA-A to PD-L1 expression levels did not exhibit a clear dependence on the type of IFNγ variants. This indicates that the expression balance of two contrastive genes follows a similar pattern regardless of whether IFNγ variants are monomeric or dimeric, with or without an NLS. In addition to HLA-A and PD-L1, the same trend of expression balance was observed for TAP1 and IRF1, suggesting that the expression balance of immune-activating and immunosuppressive genes elucidated here is not limited to HLA-A and PD-L1. These findings were obtained as a result of gene expression being measured over a wide concentration range of 0.001–1000 nM, a unique approach in this study.

**Figure 6.**
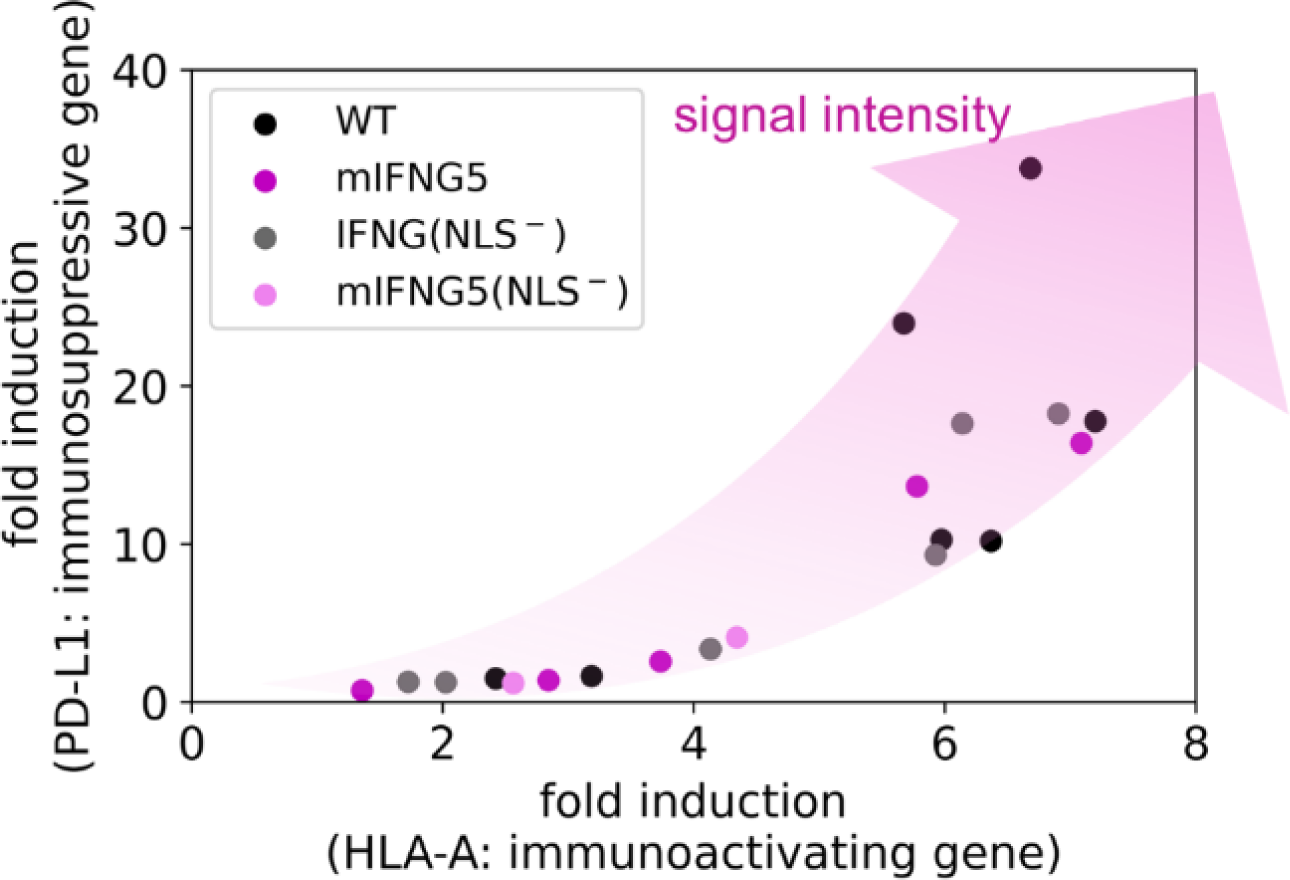
Relationship between immunostimulatory and immunosuppressive gene expression upon stimulation with IFNγ variants. The expression balance of two contrastive genes did not depend on the structural formats of the ligands but rather on their intracellular signal intensity.

Our findings suggest that the switch between immune activation and immune suppression is regulated by intracellular signal intensity rather than by the structure of IFNγ. Initially, we hypothesized that the structural formats of the ligand, such as the domain-swapped structure, the number of epitopes, and the presence of an NLS, might be implicated in the switch between immune activation and immune suppression. These postulates were based on a previous study by Mendoza et al., who found that a single-epitope IFNγ variant, GIFN2, exhibited reduced immunosuppressive function while maintaining immunostimulatory function.^16^ Although structural format certainly affects intracellular signal intensity, inferring regulation of gene expression by structural format may not be appropriate. Since the concentration-dependent induction of PD-L1 expression by GIFN2 exhibited a similar tendency,^16^ the results of Mendoza et al. may be interpreted from the viewpoint of intracellular signal intensity.

In this study, we successfully achieved the monomerization of IFNγ. The domain-swapped dimer with a large subunit interface was transformed into its monomeric form through a two-step approach involving linker insertion and substitution of hydrophobic amino acids at the subunit interface. The resulting mIFNG5 was confirmed to be monomeric and active in assays of receptor binding, STAT1 phosphorylation, and downstream gene expression. Subsequent cell-based analysis of different IFNγ variants, including mIFNG5, demonstrated that IFNγ-induced gene expression exhibits a common intracellular signal intensity-dependent pattern. Our findings suggest that molecules which bind tightly to receptors, but exhibit weak intracellular signaling, hold potential as therapeutic agents for cancer immunotherapy.

## METHOD DETAILS

### IFNγ expression vector

The sequence of WT used in this study was derived from previous studies (MQDPYVKEAENLKKYFNAGHSDVADNGTLFLGILKNWKEESDRKIMQSQIVSFYFKLFKNFKDDQSIQKSVETIKEDMNVKFFNSNKKKRDDFEKLTNYSVTDLNVQRKAIHELIQVMAELSPAAKTGKRKRSQAAAHHHHHHHH).^16,42^ The codons have been optimized for expression in *E. coli*. The codons S85 and N86 were designed as the recognition sequence (TTCGAA) of *Bsp*T104 I (Takara Bio Inc., Shiga, Japan) for further modification of the linker insertion. The synthetic gene was inserted into the *Nde* I/*Xho* I sites of the pET-24b(+) vector. Designed expression vectors were purchased from GenScript (New Jersey, USA).

### Linker insertion

The synthetic gene for a GS linker (NSNGGGGSGGGGSGGNSN) with a *Bsp*T104 I recognition sequence at its termini was purchased from Eurofins Japan (Tokyo, Japan). The GS linker synthetic gene and the WT expression vector were treated with *Bsp*T104 I and dephosphorylated using alkaline phosphatase (Toyobo Co., Ltd., Osaka, Japan). The dephosphorylated genes were mixed, and ligation was performed using a Ligation-Convenience Kit (Nippon Gene Co., Ltd., Toyama, Japan). The obtained plasmid was transformed into *E. coli* DH5α (Takara Bio Inc.) to obtain the plasmid for the linker-inserted variant.

### Vector modification

Linker extensions, deletions, and point mutations were performed as follows. Primers were designed using a PrimeSTAR Mutagenesis Basal Kit (Takara Bio, Inc.) according to the manufacturer’s protocol. The primers and template plasmids were mixed and amplified via PCR using KOD-Plus (Toyobo Co., Ltd.). The PCR product was processed with *dpn* I (Takara Bio Inc.) to digest the template plasmid and then transformed into *E. coli* DH5α to obtain the modified plasmid.

### Protein expression and purification

Expression of WT and its variants was carried out using *E. coli* BL21 (DE3) (Merck Millipore, Massachusetts, USA) transformed with the corresponding expression plasmid. Bacterial cells were cultivated in 2x YT medium (16 g/L tryptone, 10 g/L yeast extract, 5.0 g/L NaCl) until an optical density of 0.6 at 600 nm (OD600) was reached. Isopropyl β-D-1-thiogalactopyranoside was then added to a final concentration of 1 mM, and the bacterial cells were further incubated for 4 h. The bacterial cells were washed with a buffer consisting of 100 mM Tris-HCl (pH 8.0), 10 mM EDTA, and 100 mM NaCl and resuspended. The suspension was then sonicated to obtain inclusion bodies. The inclusion bodies were washed with a buffer consisting of 100 mM Tris-HCl (pH 8.0), 1.0 mM EDTA, and 1.0 M NaCl and solubilized in 6.0 M GdnHCl and 20 mM Tris-HCl (pH 8.0). The solubilized mixture was filtered through a 0.45 um filter and applied onto a His GraviTrap (Cytiva, Inc., Massachusetts, US) equilibrated with a buffer consisting of 6.0 M GdnHCl, 20 mM Tris-HCl (pH 8.0), and 20 mM imidazole, then eluted with a buffer consisting of 6.0 M GdnHCl, 20 mM Tris-HCl (pH 8.0), and 300 mM imidazole. The purified fraction was refolded via 200-fold dilution in 20 mM sodium acetate buffer (pH 5.0). The refolded solution was concentrated by TFF using a Vivaflow 50R Crossflow Cassette (Sartorius, Göttingen, Germany) and applied onto a preparative size-exclusion chromatography column (Superdex 75 10/300 GL, Cytiva, Inc.) equilibrated in 100 mM sodium acetate buffer (pH 5.0) and eluted with the same buffer. The purity of each sample was verified by analytical SEC. Each sample was confirmed as a single peak.

### Analytical ultracentrifugation (AUC)

Proteins were dissolved in 100 mM sodium acetate buffer (pH 5.0). The experiments were performed using a ProteomeLab XL-A (Beckman Coulter, Inc., Brea, California, USA) at a rotation speed of 50000 rpm. The absorbance at 280 nm was measured using a double-sector cell with a 12 mm path length. The data were collected for 15 h. Sedimentation coefficients of the proteins in water at 20 °C (*s20,w*) were obtained from analyses of the measured data using the SEDFIT program (version 16.36, 2016, Schuck)^43^ and appropriate corrections. The partial specific volume of proteins, solvent viscosity, and density used for AUC analyses were calculated using the SEDNTERP program (version 3.0.4). The continuous sedimentation coefficient distribution *c*(*s*) model^44^ was selected, and the c(*s*), molecular mass, friction ratio (*f*/*f*_0_), and hydration radius (*R*_H_) were calculated. The initial settings of *f*/*f*_0_, sedimentation coefficient analysis range, and resolution were set to 1.2, 0–15 S, and 200 s-values, respectively. Fitting was performed until the root-mean-square deviation was within 1% of the OD_280_, and the residual bitmaps were evaluated through visual inspection.

### Analytical size-exclusion chromatography (SEC)

Analytical SEC was performed on a Prominence HPLC system (Shimadzu Co., Kyoto, Japan), using a TSKgel G2000SWXL column (300 mm × 7.8 mm; Tosoh Co., Ltd, Tokyo, Japan). Aliquots of the protein solution (10 μL) were loaded onto a column equilibrated in 100 mM sodium acetate buffer consisting of 300 mM sodium chloride (pH 5.0) and eluted with the same buffer at a flow rate of 1.0 mL/min. The ultraviolet absorbance of the eluate was monitored at 280 nm.

### Evaluation of aggregation propensity under neutral conditions

Proteins were diluted in 100 mM sodium acetate buffer (pH 5.0) to an absorbance of 0.6 at 280 nm with a 10 mm path length (A280). To neutralize the solution to pH 7.5, 25 µL of 1.0 M Tris (pH 11.1) was added to 200 µL of protein solution, followed by the measurement of A280 after filtration through a 0.45 µm pore size filter (Check 1). After being left to stand at 4 °C for one day, A280 was measured again following filtration through a 0.45 µm pore size filter (Check 2). For samples without aggregation, the solution was further left to stand at room temperature (20-25 °C) for one day, and A280 was measured after filtration through a 0.45 µm pore size filter (Check 3). Samples showing a decrease in A280 at each checkpoint were considered aggregated.

### Epitope Binning

The epitope binning assay was conducted using the Octet RED 96 system (Sartorius). Anti-IFNγ antibody (ab180547, Abcam, Cambridge, UK) was immobilized on Octet Amine Reactive 2nd Generation (AR2G) Biosensors (Sartorius) at a concentration of 200 nM in a 10 mM sodium acetate buffer (pH 5.0) via a standard EDC-catalyzed amide bond formation. The analyte and anti-IFNγ antibody were diluted to a concentration of 50 nM and 100 nM, respectively, using HBS buffer (10 mM HEPES, pH 7.4, 150 mM NaCl, 3.4 mM EDTA, and 0.05% Tween20). The antibody-immobilized sensor was immersed in the anti-IFNγ antibody solution for 600 s to verify the absence of interaction between the antibodies. The analyte was then reacted by immersion in the analyte solution for 600 s, followed by immersion in the HBS buffer for 300 s. The sensor was then immersed in the anti-IFNγ antibody solution for 600 s and evaluated for the presence of a signal.

### Receptor binding assay

Kinetic assays were performed using the Octet RED 96 system. Analyses were conducted under 1000 rpm shaking at 30 °C. IFNγR1 (ab235874, Abcam) was immobilized on AR2G Biosensors at a concentration of 20 μg/mL in a 10mM sodium acetate buffer (pH 5.0) via a standard EDC-catalyzed amide bond formation. The analyte was diluted to various concentrations using HBS buffer. Biosensor-analyte association was measured by immersing the biosensor in analyte solution for 400 s, and dissociation was measured by immersing the biosensor in HBS for 800 s. The negative control, protein A, was purchased from Nacalai Tesque, Inc. (29435-14, Kyoto, Japan).

### Phosphorylated STAT1 (pSTAT1) assay

The AlphaLISA pSTAT1 assay was performed using the AlphaLISA SureFire Ultra p-STAT1 (Tyr701) assay kit (PerkinElmer, Inc., Waltham, MA, USA). HeLa cells (CCL-2; ATCC, Manassas, Virginia, USA) were seeded into a 96-well plate at a density of 40,000 cells/well and cultured in 10% BSA-supplemented RPMI-1640 medium (Wako Pure Chemical Industries, Ltd., Osaka, Japan) at 37 °C overnight. The medium was removed, and various concentrations of WT and variants in BSA-free RPMI-1640 medium were added to the cells which were then incubated at 37 °C. The signal was measured using an EnSpire Alpha (PerkinElmer, Inc.) according to the manufacturer’s protocol. To minimize sample degradation, protein solutions were frozen and stored at -80 °C immediately after preparation. The aliquoted samples were thawed and used immediately prior to the pSTAT1 assay. No effect of freezing and thawing on sample quality was confirmed with analytical SEC using WT and mIFNG5.

### Quantitative reverse transcription PCR (RT-qPCR)

Hela cells were seeded into a 6-well plate at a cell density of 3.6×10^5^ cells/well and incubated overnight at 37 °C in 10% BSA-supplemented RPMI-1640 medium. Various concentrations of WT and variants were added to the cells which were then incubated at 37 °C for 48 h. After harvesting 1×10^6^ cells, RNA was extracted using a High Pure RNA Isolation Kit (Roche, Basel, Switzerland). RT-qPCR for 18S rRNA, HLA-A, and PD-L1 was performed on 2.5 ng of RNA using the LightCycler EvoScript RNA SYBR Green I Master (Roche) and the LightCycler 480 Instrument II (Roche), according to the manufacturer’s protocol. Primers for 18S rRNA (fwd 5′GTAACCCGTTGAACCCCATT3′, rev 5′CCATCCAATCGGTAGTAGCG3′), HLA-A (fwd 5′AGATACACCTGCCATGTGCAGC3′, rev 5′GATCACAGCTCCAAGGAGAACC3′), and PD-L1 (fwd 5′TGGCATTTGCTGAACGCATTT3′, rev 5′TGCAGCCAGGTCTAATTGTTTT3′) were purchased from Eurofins Japan. The amount of RNA was calculated from the *C*_p_ value obtained via the 2nd derivative max method using the LightCycler 480 software version 1.5.1.62 SP3. The relative expression of HLA-A and PD-L1 was calculated by dividing the amount of HLA-A and PD-L1 transcripts by that of 18S rRNA, the reference gene. Fold induction was calculated by dividing the relative expression of each sample by that of the unstimulated negative control (NC).

### Western blotting

Hela cells were seeded into a 6-well plate at a cell density of 3.6×10^5^ cells/well and incubated overnight at 37 °C in a 10% BSA-supplemented RPMI-1640 medium. Various concentrations of WT protein and IFNγ variants were added to the cells, which were then incubated at 37 °C for 48 h. The cells were lysed with lysis buffer (ALSU-LB-10mL, PerkinElmer, Inc.) and shaken at 350 rpm for 10 min at room temperature (20-25 °C). The lysates were separated using SDS-PAGE and transferred onto nitrocellulose membranes. The membranes were blocked in blocking buffer (PBS containing 5.0% skim milk and 0.05% Tween 20) by shaking for 1 h at room temperature. Subsequently, appropriate primary antibodies were added to the blocking buffer, and the membranes were incubated overnight at 4 °C. The membranes were washed with PBS containing 0.05% Tween 20, followed by the addition of appropriate horseradish peroxidase (HRP)-conjugated secondary antibodies in blocking buffer and incubation for 1 h at room temperature. The membranes were washed with PBS containing 0.05% Tween 20. Luminata Crescendo Western HRP Substrate (WBLUR0100, Merck Millipore) was added to the membranes, which were then visualized using Fusion FX (Vilber, Collégien, France). This study was performed using the following antibodies: anti-HLA-A (ab52922, Abcam), anti-PD-L1 (ab205921, Abcam), anti-β-actin (sc-47778, Santa Cruz Biotechnology, Texas, USA), anti-rabbit IgG (HRP) (ab205718, Abcam), and anti-mouse IgG (HRP) (#32430, Thermo Fisher Scientific, Inc., Massachusetts, USA).

## ACKNOWLEDGMENTS

We thank Dr. Risa Shibuya for the guidance with the experiments and data analyses. We also thank Drs. Yukako Senga, Hideki Watanabe, and Shota Shiga for their advice regarding the experiments.

## AUTHOR CONTRIBUTIONS

Y.G. designed the study, performed the experiments, analyzed the data, and wrote the initial draft of the manuscript. T.M. advised on the experiments and data interpretation. S.H. designed the study, provided guidance, and edited the manuscript. All authors contributed to the data interpretation and critically reviewed the manuscript. The final version of the manuscript has been approved by all authors.

