## Supporting Information for "Designing monomeric IFNγ: the significance of domain-swapped dimer structure in IFNγ immune responses"

TABLE OF CONTENTS

Supporting Tables

Table S1. Comparison of theoretical (calculated) and observed MW and $S_{20,w}$ values, related to Figure 3A, B, C

Table S2. Equilibrium dissociation constant ($K_d$) calculated from sensorgram association and dissociation signals, related to Figure 4B

Supporting Figures

Figure S1. Reproducibility of RT-qPCR measurements, related to Figure 5F, G

Figure S2. Phylogenetic tree on IFNγ in various species and their structures

Figure S3. Multiple sequence alignment of mammalian IFNγ

**Table S1. Comparison of theoretical (calculated) and observed MW and $S_{20,w}$ values, related to Figure 3A, B, C**

| | theoretical MW | observed MW | calculated $s_{20,w}$ | observed $s_{20,w}$ | condition |
|---|---|---|---|---|---|
| WT (dimer) | 34029 | 33947 | 2.31 | 2.38 | a* |
| mIFNG1 | 17873 | $2\times10^5$-$5\times10^5$ | 1.63 | 8-20 | a |
| WT (dimer) | 34029 | 33555 | 2.31 | 2.44 | b** |
| mIFNG5 | 17850 | 19106 | 1.65 | 1.64 | b |

*a: Proteins were prepared in 100 mM sodium acetate buffer (pH 5.0) with an absorbance of 0.8 at 280 nm (A280). Experiments were performed at 20 °C and 50000 rpm.

**b: Proteins were prepared in 100 mM sodium acetate buffer (pH 5.0) with an A280 of 0.2. Experiments were performed at 10 °C and 50000 rpm.

**Table S2. Equilibrium dissociation constant ($K_d$) calculated from sensorgram association and dissociation signals, related to Figure 4B**

| | $K_{d,1}$ | $K_{d,2}$ |
|---|---|---|
| WT (reported)* | $4.50\times10^{-9}$ | |
| WT | $7.80 \pm 2.14\times10^{-9}$ | $1.83 \pm 0.66\times10^{-10}$ |
| mIFNG5 | $2.43 \pm 1.14\times10^{-8}$ | $4.11 \pm 1.41\times10^{-10}$ |

Signals were fitted using a 2:1 fitting model with the system software data analysis (version 7.0), and the $K_d$ was calculated. Data represent the mean ± standard deviation (STD) of $K_d$ and $K_{d,2}$ from three independent experiments (n=3). *Data obtained from the literature[40].

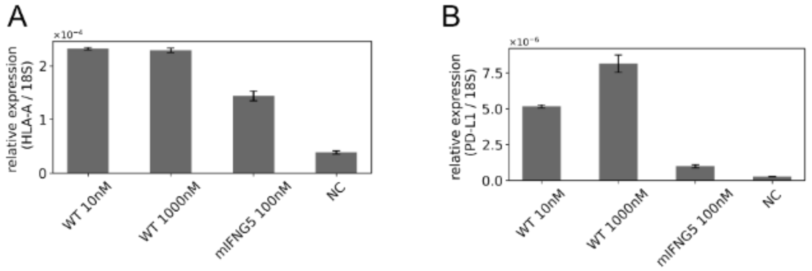


**Figure S1. Reproducibility of RT-qPCR measurements, related to Figure 5F, G**

(A, B) Reproducibility of RT-qPCR measurements. Relative expression of target genes HLA-A (A) and PD-L1 (B) under several conditions. Data represent the mean relative expression ± standard error of the mean (SEM) of n=3 biological replicates.

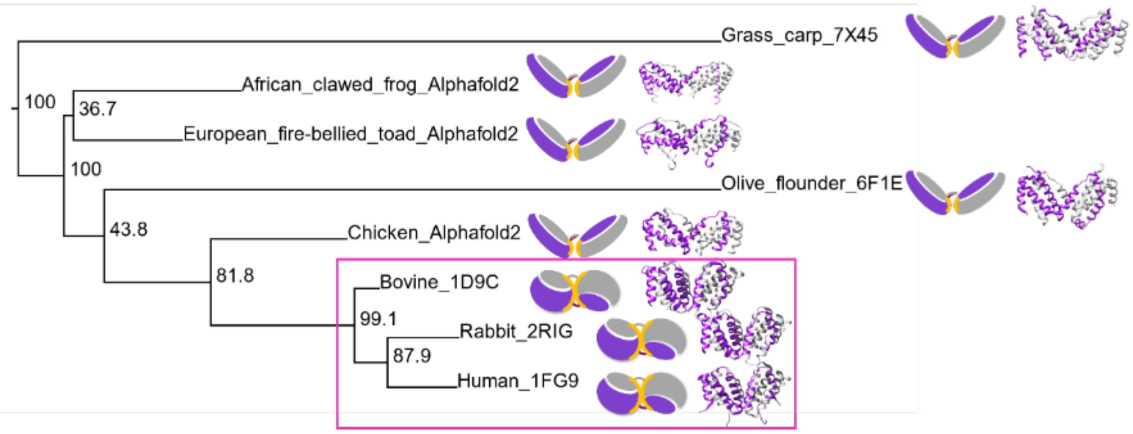


**Figure S2. Phylogenetic tree of IFNγ in various species and their structures**

The phylogenetic tree was constructed from multiple sequence alignments of IFNγ in various species using CLUSTALW. Ribbon models and simplified structures are shown in the phylogenetic tree. The structures of IFNγ of the species whose structures have been determined were taken from PDB (PDB ID is shown to the right of the species name). Predicted structures of IFNγ in species for which the structure has not been determined were generated by Alphafold2.

```
human_1FG9_    MQDPYVKEAENLKKYFNAGHSDVADNGTLFLGILKNWKEESDRKIMQSQIVSF
rabbit_2RIG_   -QDTLTRETEHLKAYLKANTSDVANGGPLFLNILRNWKEESDNKIIQSQIVSF
bovine_1D9C_   -QGQFFREIENLKEYFNASSPDVAKGGPLFSEILKNWKDESDKKIIQSQIVSF

human_1FG9_    YFKLFKNFKDDQSIQKSVETIKEDMNVKFFNSNKKKRDDFEKLTNYSVTDLNV
rabbit_2RIG_   YFKLFDNLKDHEVIKKSMESIKEDIFVKFFNSNLTKMDDFQNLTRISVDDRLV
bovine_1D9C_   YFKLFENLKDNQVIQRSMDIIKQDMFQKFLNGSSEKLEDFKKLIQIPVDDLQI

human_1FG9_    QRKAIHELIQVMAELSPAAKTGKRKRSQ-----------
rabbit_2RIG_   QRKAVSELSNVLNFLSPKSNLKKRKRSQTLFRGRRASKY
bovine_1D9C_   QRKAINELIKVMNDLS-----------------------
```

**Figure S3. Multiple sequence alignment of mammalian IFNγ**

Multiple sequence alignment of three mammalian IFNγ with determined structures, human (PDB: 1FG9), rabbit (PDB: 2RIG, aligned score 61.9, RMSD (Cα) 2.77 with human), and bovine (PDB: 1D9C, aligned score 60.3, RMSD (Cα) 1.04 with human) IFNγ, using CLUSTALW. Highlighted in yellow are the positions of hydrophobic amino acid residues that are the focus of this study.
